# Disentangling *Respiratory Phase-Dependent* and *Anticipatory* Cardiac Deceleration in a Visual Perception Task

**DOI:** 10.1101/2025.08.15.670501

**Authors:** Ege Kingir, Sukanya Chakraborty, Caspar M. Schwiedrzik, Melanie Wilke

**Affiliations:** Department of Cognitive Neurology, Heart and Brain Center Goettingen, University Medical Center, Robert-Koch-Str. 42, Goettingen, 37075, Germany; Cognitive Neurology Group, German Primate Center, Cognitive Neuroscience Laboratory, Leibniz Institute for Primate Research, Kellnerweg 4, Goettingen, 37077, Germany; Neural Circuits and Cognition Lab, European Neuroscience Institute Goettingen – A Joint Initiative of the University Medical Center Goettingen and the Max Planck Institute for Multidisciplinary Sciences, Grisebachstraße 5, 37077 Goettingen, Germany; Perception and Plasticity Group, German Primate Center – Leibniz Institute for Primate Research, Kellnerweg 4, 37077 Göttingen, Germany; Cognitive Neurobiology, Research Center One Health Ruhr, University Alliance Ruhr, Faculty of Biology and Biotechnology, Ruhr-University Bochum, Universitätsstraße 150, 44801 Bochum, Germany

**Author notes:** corresponding author: Corresponding author: Melanie Wilke. first author (Ege Kingir). shared senior authors (Caspar M. Schwiedrzik, Melanie Wilke).

**Keywords:** anticipatory cardiac deceleration, heart rate variability, respiration, respiratory sinus arrhythmia, visual perception

## Abstract

The heart does not beat like a metronome: varying parasympathetic input to the heart leads to constant heart rate variability. Vagal cardiomotor neuron activity is coupled to the respiratory cycle, leading to Respiratory Sinus Arrhythmia (RSA), a permanent oscillation of heart rate synchronized to respiration. Heart rate also temporarily decelerates in specific conditions such as in freezing due to perceived threat, or anticipation of a salient stimulus. Anticipatory Cardiac Deceleration (ACD) is observed consistently in anticipation of a stimulus in perceptual tasks, but its relationship with perceptual performance is debated. Previous quantifications of ACD neglect ongoing heart rate oscillations due to RSA, which may have led to inconsistencies in the ACD-related analyses across studies. Here, we suggest a novel approach to estimate trial-averaged RSA amplitude and respiratory phase-independent cardiac deceleration simultaneously, and apply it to an EEG-ECG dataset from a visual detection task. While the total ACD was not associated with perception, dissociating RSA-based and non-respiratory cardiac modulations revealed that they show opposing effects on perceptual performance. Additionally, we found that participants with higher ACD amplitudes also displayed larger Visual Awareness Negativity potentials, further supporting a contribution of ACD to visual perception.

**Impact Statement:** We present a novel analysis method to quantify task-related, anticipatory cardiac deceleration which takes tonic heart rate oscillations due to respiratory sinus arrhythmia into account. Our results add to previous research on the relationship between cardiac deceleration and perception by simultaneously characterizing and dissociating respiratory and non-respiratory heart rate modulations during stimulus anticipation.

## 1. INTRODUCTION

The brain constantly communicates with the rest of the body as we sleep, rest, or perform any physical or mental task. Ascending neural pathways from the visceral organs such as the heart and lungs keep the brain informed about the bodily state of an organism (Azzalini et al., 2019). The cortical and subcortical brain regions that receive and process these interoceptive inputs are collectively referred to as the ‘Central Autonomic Network’ (CAN) (Shaffer et al., 2014; Sklerov et al., 2019). Higher-order cortical regions considered to be a part of the CAN include –but are not limited to– the insular cortex, cingulate cortex, and the amygdala (Ferraro et al., 2022; Shaffer et al., 2014). The CAN receives interoceptive information by afferent neurons relayed on the medulla oblongata of the brainstem, and integrates it with the exteroceptive information (Benarroch, 1993). The output of the CAN, in turn, mainly controls the balance between the parasympathetic and the sympathetic branches of the Autonomic Nervous System (ANS), thus producing integrated autonomic and behavioral responses in accordance with the environmental demands and the visceral state of the organism (Benarroch, 1993; Shaffer et al., 2014). These efferent pathways, mainly implemented through several branches of the vagus nerve, control both cardiac and respiratory outputs (Benarroch, 1993; Sevoz-Couche & Laborde, 2022).

As a result of the ever-changing vagal tone on the heart, the time between successive heartbeats fluctuates constantly. The amount of this variance in a given time window is called the Heart Rate Variability (HRV) (Shaffer & Ginsberg, 2017). Since the output from the CAN is considered to be the main source of variance in the interbeat intervals (IBIs), HRV is also used as an index for how much an organism can flexibly adjust its autonomic functions (e.g., heart rate) in response to varying environmental or cognitive demands (Skora et al., 2022; Thayer & Lane, 2000). However; HRV is caused by multiple factors as we will introduce below, and traditional HRV measurements require continuous recordings in the range of minutes (Shaffer & Ginsberg, 2017). Therefore, it has been difficult for previous studies on HRV to extract and dissociate individual contributors to the total variability of heart rate, especially in short time windows.

Several conditions such as observing a distant threat, anticipation of a task-relevant signal, or receiving an error-related signal induce a robust temporary cardiac deceleration (Skora et al., 2022; Thayer & Lane, 2000). All of these conditions require the organism to selectively attend to the relevant event (Thayer & Lane, 2000). The amount of temporary cardiac deceleration under such circumstances is associated with behavioral flexibility (Thayer & Lane, 2000). It has also been hypothesized that the increased parasympathetic input to the heart allows the brain to assign lower levels of precision to the interoceptive signals, thereby optimizing the processing of exteroceptive signals with a precision weighting mechanism (Skora et al., 2022). The temporary cardiac deceleration that occurs in anticipation of a task-relevant stimulus is called Anticipatory Cardiac Deceleration (ACD). In the field of experimental psychophysiology, ACD is observed mainly between the cue and the target signal in “Go/No-Go” tasks or similar response timing tasks (Aprile et al., 2024; Ribeiro & Castelo-Branco, 2019). A recent study showed that ACD amplitudes are positively correlated with accuracy and faster response timings in targeted actions (Alam et al., 2023). However; preparation for a motor response is not strictly required for ACD (Steinhauer et al., 1992). The phenomenon is attributed to anticipatory attention, such that in any task where the timing of a relevant target event is predictable to some extent, ACD is observed in the last few seconds prior to target onset (Somsen, Jennings, & Van der Molen, 2004; Steinhauer et al., 1992).

ACD in trial-based paradigms is typically derived from the increasing time between consecutive R-peaks on the ECG signal during the anticipation period, and it is assumed that this measure directly corresponds to the short-term regulation of the heart rate due to anticipation (Ribeiro & Castelo-Branco, 2019; Skora et al., 2022). However; temporary environmental or cognitive demands are not the only modulators of heart rate. Instead, the heart rate constantly oscillates at the frequency corresponding to the respiratory cycle, which is known as ‘Respiratory Sinus Arrhythmia’ (RSA) (Berntson et al., 1993), or recently introduced as ‘Respiratory HRV’ (Menuet et al., 2025). A central network at the brainstem inhibits vagal cardiomotor neurons during inhalation, and this inhibition is lifted during exhalation, causing the heart rate to increase during inhalation and to decrease during exhalation (Berntson et al., 1993; Elstad et al., 2018; Shaffer et al., 2014). Although its amplitude is found to be slightly decreasing during task performance (Overbeek et al., 2014), RSA is a prominent phenomenon at all times. We suggest that previous quantifications of ACD include RSA-based modulations of the heart rate, alongside the cognitive and non-respiratory parasympathetic break due to anticipation. Since RSA is a permanent source of HRV which may drop in magnitude during task performance while ACD is prominent only in certain task-related conditions; RSA and non-respiratory ACD might also be differentially related to the behavioral outcome in the task of interest.

Disentangling RSA-based heart rate modulations while quantifying ACD might become particularly useful in cases where ACD is observed, but the findings about its relevance to the behavioral task are inconsistent. For instance, several studies reported prominent ACD in perceptual tasks, in which participants were asked to detect low-intensity stimuli at their detection thresholds (Grund et al., 2022; Motyka et al., 2019; Park et al., 2014). Unlike in tasks with high response timing pressure (Alam et al., 2023; Ribeiro & Castelo-Branco, 2019), the relevance of ACD to the behavioral outcome in these perceptual tasks is not entirely clear. Despite the prominently observed ACD in all studies cited above, the amplitude of ACD is found to be similar in trials with successful and unsuccessful perception of a target stimulus (Grund et al., 2022; Park et al., 2014), with one exception showing an increased ACD in trials with successful somatosensory stimulus detection (Motyka et al., 2019). It is known that ACD occurs due to a parasympathetic brake upon recognition of specific environmental demands (Roelofs, 2017; Skora et al., 2022), which implies that the relevant heart rate modulations in the aforementioned paradigms correspond to the top-down parasympathetic input on the heart, initiated by the cognitive task of orienting to the anticipated event. However, whether this parasympathetic “brake” facilitates sensory perception –and how it affects stimulus-evoked brain potentials– remains unresolved.

Theoretically, the differing results in the perceptual studies mentioned above can be explained by a combination of several factors such as the length and variability of the anticipatory time window, the sensory modality of choice (visual vs. somatosensory), and potential demographic differences between the participant samples (Grund et al., 2022; Motyka et al., 2019; Park et al., 2014). Yet another explanation for the inconsistent findings could be potential differences in respiratory dynamics, altering the net effect of RSA on heart rate. Previous studies suggest that participants align their respiratory phases differently based on the specific demands of the task at hand. For instance, a recent study found a respiratory phase alignment to late exhalation at the moment of voluntary motor action (Park et al., 2020), while another study found an alignment to exhalation onset at the moment of somatosensory stimulus presentation (Grund et al., 2022). Interestingly, another study suggests that visual perception threshold decreases in trials when the stimulus onset occurs during early phases of inhalation (Kluger et al., 2021). In addition, the respiratory phase appears to be coupled with pupil size oscillations, suggesting a relationship between arousal and respiration (Kluger et al., 2024). Therefore, it is likely that participants align their respiratory phases to different parts of the respiratory cycle in different tasks. This would cause the heart rate to be modified in different directions due to respiratory phase-dependent effect of RSA, potentially explaining the inconsistencies regarding the correlations between ACD amplitude and sensory perception.

The onset of ACD with respect to the stimulus onset also differs between studies that quantify ACD at multiple time points within the anticipatory window, some showing an earlier onset of cardiac deceleration (Motyka et al., 2019; Park et al., 2014), and another study showing a deceleration only on the last IBI before stimulus onset (Grund et al., 2022). These differences could also be explained by differences in the respiratory dynamics. For instance, if there is a group-level respiratory phase alignment to exhalation since the beginning of the anticipatory window, RSA would also support cardiac deceleration. Whereas if the respiratory phase is mostly aligned to inhalation, RSA would act to accelerate the heart rate while the non-respiratory parasympathetic break would act in the other direction. Therefore, quantifying trial-based respiratory dynamics and RSA amplitudes could help to disentangle respiratory (RSA-based) and non-respiratory (cognitive, top-down parasympathetic input irrespective of the respiratory phase) heart rate modulations, potentially reconciling ACD-related findings across studies.

If ACD is truly contributing to better performance in perceptual tasks, this could be mediated by an influence of ACD on sensory evoked potentials in the brain. For visual stimuli, an evoked potential known as Visual Awareness Negativity (VAN) is associated with conscious perception of the stimulus (Eklund & Wiens, 2018; Koivisto & Grassini, 2016). VAN is characterized by a negative deflection in the occipital and posterior-temporal recording sites, around 200 ms after the stimulus, and it is mostly absent if the participant does not become aware of the visual stimulus (Koivisto & Grassini, 2016; Koivisto & Revonsuo, 2003, 2010). In relation to its postulated role in anticipatory attention and precision adjustment to favor the processing of task-relevant stimuli (Skora et al., 2022; Somsen, Jennings, & Van der Molen, 2004; Steinhauer et al., 1992), ACD –and particularly the non-respiratory component of cardiac deceleration– might also enhance the VAN amplitudes, leading to a higher chance of detecting a threshold-level stimulus. Dissociating the respiratory and non-respiratory cardiac deceleration during stimulus anticipation enables us to test these ideas.

There are several established methods to estimate RSA. Most common estimators are the power of the respiratory frequency band (0.12-0.40 Hz or 0.15-0.40 Hz) from the IBI time-courses (Berntson et al., 1993) and the average difference between the longest and shortest IBIs within singular respiratory cycles (Lewis et al., 2012). The Porges-Bohrer method builds upon the frequency-power based estimation method: it interpolates the R-R interval time-series and extracts its power in the respiratory band from sliding windows of typically 15-60 seconds (Lewis et al., 2012). There have even been modifications to the Porges-Bohrer method to estimate second-by-second fluctuations in RSA (Abney et al., 2021; Fisher et al., 2016), but these methods also rely on sliding windows of at least a couple of seconds, and are thus incompatible with most trial-based approaches as in perceptual experiments. In addition, these methods only quantify the respiratory phase-dependent effect of RSA, omitting the non-respiratory modulations of the heart rate.

It is difficult to dissociate RSA from ACD on a trial-by-trial basis, especially since both phenomena are quantified by the change in heart rate. However, one can take advantage of the idea that while RSA is a respiratory phase-dependent modulator of the heart rate, the effect of ACD should be similar at all respiratory phases. Here, we suggest a novel approach to estimate trial-averaged RSA amplitudes from non-overlapping data points. We demonstrate that this method can also disentangle the relative contributions of respiratory phase-dependent (RSA) and phase-independent heart rate modulations that may co-occur during specific environmental or cognitive demands such as in anticipation of a target event. Specifically, we show that sinusoidal modeling of the trial-averaged changes in heart rate as a function of the respiratory phase can simultaneously reveal RSA amplitude (by the amplitude of the sinusoidal fit), and the non-respiratory cardiac deceleration (by the vertical offset of the sinusoidal fit). We also investigate the relationship between these cardiac deceleration parameters and both the perceptual performance and the cortical stimulus-evoked potentials in a threshold-level visual detection task.

## 2. METHODS

### 2.1. Participants

A total of 28 healthy adult participants with normal or corrected-to-normal vision participated. 1 participant was excluded due to a large proportion of false alarms in the stimulus-absent trials (more than a median of 1 false alarm per 9 stimulus-absent trials in a block), and 4 more participants were excluded due to low quality respiration data. In total, 8 male and 15 female participants were included in the data analysis, aged between 22 and 34 (mean ± SD: 26.49 ± 3.18). The study was conducted in accordance with the ethical guidelines of the Declaration of Helsinki. The experimental procedure was approved by the ethics committee of the University Medicine Göttingen. All subjects gave written informed consent prior to the start of the experiment and were paid for their participation.

### 2.2. Experimental Recordings

#### 2.2.1. ECG acquisition

For the ECG setup, 4 disposable cup electrode cables (EasyCap GmbH, Wörthsee, Germany) were used with 3M Red Dot Adhesive Electrodes. Two electrodes were placed under the clavicle bilaterally and two electrodes above the pelvic bone bilaterally to form two cross recording channels for heart muscle electrical signals and one reference electrode positioned on the back. The data were recorded at a sampling rate of 1000 Hz.

#### 2.2.2. Respiration acquisition

For measuring respiratory patterns, the Brain Products Respiration Belt was used (Breathing belt, BB), placed over the participant’s chest (primarily for males) or abdomen (primarily for females) ensuring that the pressure-sensitive cushion sensor was positioned at the point of maximal inflation during inspiration. ExG amplifiers and AUX adapters from BrainVision (Brain Products GmbH, Gilching, Germany) were used for ECG and respiration recordings. The data were recorded at a sampling rate of 1000 Hz.

#### 2.2.3. EEG acquisition

EEG activity was recorded using 64 channels in a 10-20 system cap (actiCAP 64-channel active system, BrainCap, BrainVision BrainAMP MR Plus or DC amplifiers, BrainVision powerpacks, BrainVision USB 2 Adapter, SyncBox; Brain Products GmbH, Gilching, Germany). Electrode impedances for EEG were kept below a maximum of 15 kΩ. The data were recorded at a sampling rate of 1000 Hz. Triggers were sent to the EEG computer via the Brain Products BrainVision SyncBox with a trigger cable connecting the stimulus presentation computer and the EEG station. EEG-ECG-BB data was recorded using the BrainVision Recorder (Brain Products GmbH, Gilching, Germany).

#### 2.2.4. Pupillometry

Eye tracking for horizontal and vertical eye positions was carried out with the Pupil Labs Eye tracking system (Pupil Labs GmbH, Berlin, Germany): wearable Pupil Core headset with two eye cameras and a scene camera (called the “world” camera in the Pupil Capture software) coupled with the Pupil Capture software. Eye cameras on the headset were physically adjusted by rotating the ball joints to get a clear image of the eyes and ensuring that the pupils were always in view and detected with high confidence (>0.8 out of 1.0 according to the Pupil Capture software). “3D pupil detection” setting was enabled for eye model estimation (to minimize the side effects of headset slippage during the experiment). Data were recorded at a sampling rate of 200 Hz, were only used for online fixation control during the experiment and not analyzed further.

#### 2.2.5. Experiment set-up

Following preliminary checking of the set-up and signals to ensure minimal noise in the raw data, participants were instructed about the task (see below) and moved to the electrically insulated, sound-proof dark participant cabin (no sources of extraneous light), where they were seated in front of a computer screen with 60 Hz refresh rate and 1920x1080-pixel resolution (BenQ XL2411T; BenQ, Taipei, Taiwan). Participants’ heads were placed on a chin rest with an eye-to-screen distance of 70 cm and fixed with an Arrington Headlock system (HeadLock Ultra Precision Head Positioner; Arrington Research, Scottsdale, AZ, USA). 10 minutes of resting-state EEG, ECG, and respiration recordings were done prior to the start of the experiment. Participants’ heads were placed on the chin rest during the resting-state recording as well, and their eyes were closed. At the end of 10-minutes, we verified that the participants did not fall asleep during the resting-state, which could have affected the cardiorespiratory interactions significantly (Stoakley et al., 2017).

### 2.3. Behavioral Task Paradigm

Participants engaged in a visual detection task. Trial sequence of the task is shown in **Fig. 1A**. The sequence was largely adapted from Park et al., 2014.

Trials started with a Fixation period, followed by a color change of the fixation cue, “warning” the participant that the target grating is about to be presented. During this anticipatory phase (and also until the end of the post-stimulus delay period), participants had to stabilize their gaze at the central fixation spot for successful completion of the trial (**see section 2.3.1.** below). Following onset of the target grating and a delay period, participant pressed one of the two assigned buttons on the button box (Current Designs Inc., Philadelphia, PA, USA) to report their perceptual decision (target perceived or not perceived). Then, they reported their decision confidence by sliding a red pointer to the left or right on a continuous scale, until the pointer reached their confidence level.

We presented the target stimulus in 87% of the trials. No stimulus was shown in the remaining trials, in order to verify that the participants attended to the task and did not produce excessive false alarms (more than a median of 1 false alarm per block). Participants performed 6-8 experimental blocks of 70 trials each.

**Figure 1:**
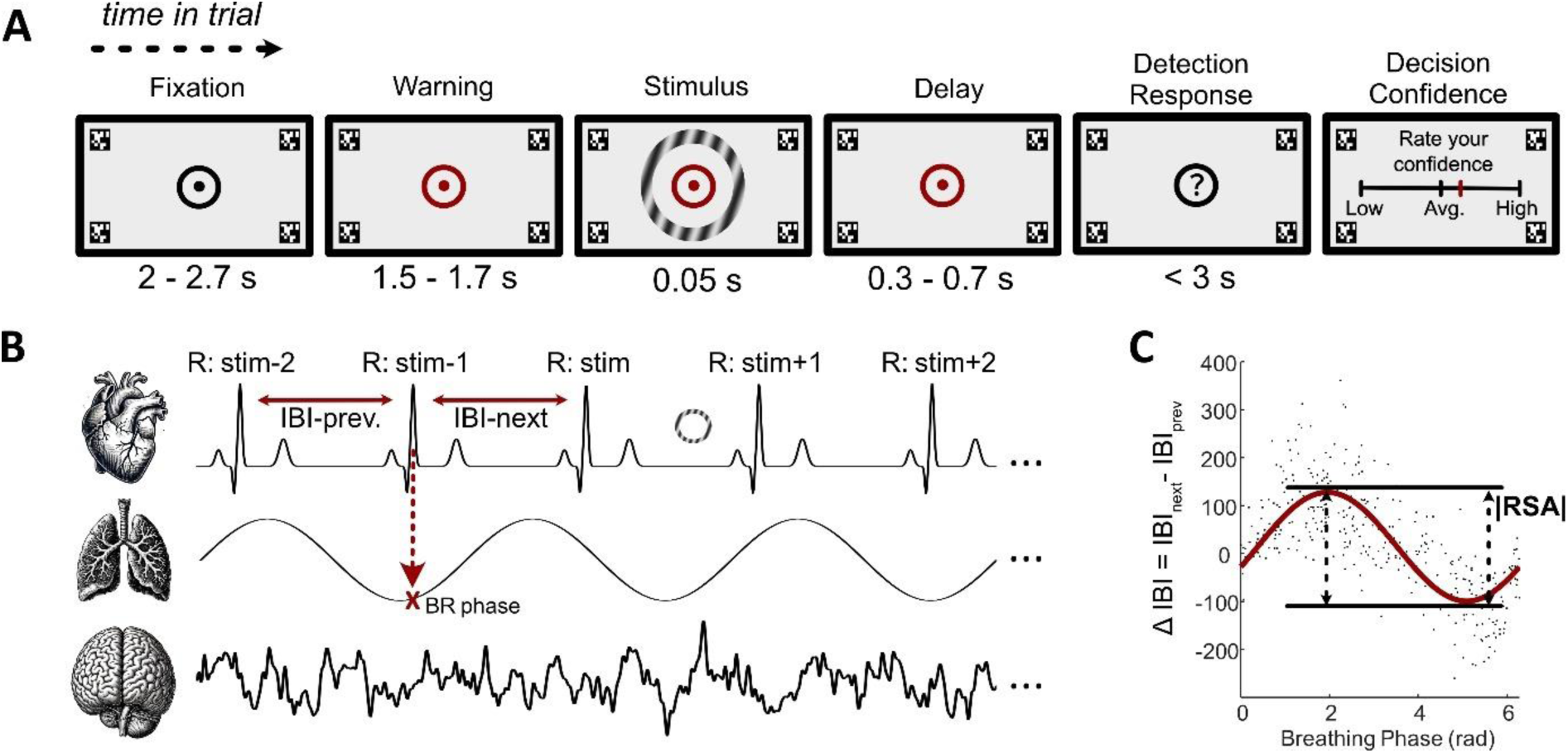
Experimental paradigm and the RSA modeling approach. **A)** Visual threshold-level stimulus detection task. The tiles represent the visual display presented to the participants during each trial. Duration ranges of each state in a trial are given below the corresponding tile. The contrast of the target grating is increased in this figure to improve its visibility to the reader. QR codes at the corners of the screen are April Tags used for the eye-tracking software to recognize the screen and compute gaze locations within this surface. **B)** Physiological recordings. ECG (top), respiration data (middle), and EEG (bottom row) were continuously recorded throughout each experimental block. During data analysis, R-peaks were organized according to their timing relative to the target stimulus onset. We refer to the last R-peak before the target onset as “R:stim”, and to the first R-peak before the target onset as “R:stim-1”, and the rest of the annotation pattern follows intuitively. **C)** Beat-by-beat, trial-averaged RSA estimation. The respiratory phase and the change in the IBI centered around a chosen R-peak (e.g., “R:stim-1”) are placed on the x- and y-axis, respectively. Each trial yields one such data point, and the overall data from the participant (scattered black dots) can be accurately summarized by a sinusoidal function (the red curve) with an amplitude, a vertical offset, and a phase shift. Amplitude of the sinusoidal function is used as the estimator for the Respiratory Sinus Arrhythmia (RSA) amplitude at that particular moment of the trial. The plot corresponds to data points from a single participant at the R-peak named “R:stim-1”.

#### 2.3.1. Eye-Tracking Calibration and Controlling for the Fixation Breaks

Each block began with an eye-tracking calibration, wherein the participant was instructed to fixate on bullseye calibration markers displayed at 5 different sequential positions on the screen in two calibration and two validation cycles to check for angular accuracy and precision of eye-tracking (automated algorithm on the Pupil Capture software v3.5.1, Pupil Labs GmbH, Germany). Once satisfactory angular accuracy and precision values were achieved (below 2.5 and 0.1 visual degrees for each participant, respectively), the fixation mark was displayed at the center of the screen with the central dot being white instead of black for 5 seconds and the average gaze position during the last 3 seconds of this period was used to estimate the mean starting gaze for each participant in each block for subsequent fixation detections.

During the experiment, ‘April Tags’ were continuously displayed on the corners of the display screen to define the target surface and acquire normalized gaze positions on the screen (see **Fig. 1A**). From the beginning of the “Fixation” to the end of “Delay” state in each trial, the normalized gaze positions on the display screen were relayed to MATLAB. This input was then used to check for fixation breaks during the experiment, in accordance with a fixation break check rule: no saccades or rapid eye movements larger than 1.5° of visual angle centered on a circular window of this radius from the subject’s center gaze location for 0.5 to 0.7 seconds. Any violations of this rule led to the trial being aborted at that state. To account for slow eye tracking drifts over trials, the center location for the participant’s gaze was updated in each “n^th^” trial by considering the weighted mean of 90% of all the center gaze locations up to and excluding the “(n – 1)^th^” and 10% of the center gaze location of the “(n – 1)^th^” trial. A blink allowance of 200 ms was also accounted for. Fixation was not controlled for during the perceptual report, confidence rating and intertrial interval states.

#### 2.3.2. Stimulus Properties

Stimuli were programmed and presented in MATLAB 2017a (The MathWorks Inc., Natick MA, USA) using Psychtoolbox-3. The central target stimulus was a grating annulus characterized by a spatial frequency of five cycles per degree of visual angle. The annulus dimensions for the inner and outer rings measured 3.0° and 3.6° of visual angles from the center of the screen, respectively. The orientation of the grating pattern for the annulus was confined to a subset of 20 possible orientations, evenly distributed between 0° and 180°, with cardinal orientations. The fixation mark consisted of a black dot with a radius of 0.15° of visual angle, encircled by a black circular ring with an inner and outer radius of 0.35° and 0.42° of visual angle respectively, positioned centrally on the screen. All visual stimuli were displayed against a grey background, presented at a viewing distance of 70 cm.

#### 2.3.3. Training and Threshold Assessment

The QUEST package under Psychtoolbox-3 in MATLAB was employed for controlling task performance rates in the visual detection experiment. Stimulus contrast was adjusted to aim for a hit rate (proportion of correct detections of the stimulus in stimulus-present trials) of 65%. To this end, each experimental session began with a short contrast calibration block (30 stimulus-present and 5 stimulus-absent trials) during which the participant performed the task exactly as described above. The stimulus contrast was varied from trial to trial using the QUEST algorithm (Watson & Pelli, 1983), a Bayesian adaptive procedure, to estimate the detection threshold from a series of psychophysical trials with a binary trial outcome (i.e., hit or miss). The threshold obtained was then used as the initial estimate of the stimulus intensity (tGuess) that is expected to result in a response rate of 65% hits in the subsequent experimental trials. Before the calibration block, the tGuess was set to a default of 0.05 log contrast value for the grating annulus stimulus intensity. Step size of contrast modulation was standardized to 0.0025, with a maximum contrast value of 0.09. This calibration block also served as the training block, and if the participant had some questions or uncertainties regarding the main experiment, the training block was repeated.

In each experimental block, the stimulus contrast was modulated consistently in the first 10 trials using the QUEST update rule to account for any changes in the detection thresholds during the self-paced breaks between blocks. Essentially, first the “QuestQuantile()” MATLAB function was utilized to provide a suggested estimate of the contrast for the next trial. This suggestion was then used to update the log contrast stimulus intensity using the “QuestUpdate()” MATLAB function accounting for the participant’s response in the current trial. After the first 10 trials, the stimulus contrast was kept constant unless the participant’s accuracy decreased to 40% or reached 80% in the preceding 10 trials. Stimulus contrast adjustment was maintained within the experimental blocks because the participants’ detection threshold can change even within a block of 8-10 minutes, which leads to many consecutive trials in which the task is impossible or too easy to perform if the stimulus contrast is kept constant throughout the block.

### 2.4. Data Preprocessing

#### 2.4.1. ECG data pre-processing

Physiological signals that are analyzed in this study are illustrated in **Fig. 1B**. R-peaks were detected after bandpass filtering the ECG data between 0.5-45 Hz, the frequency range that mainly contains the QRS complex (Martis et al., 2013; Sharma & Pachori, 2018). IBIs were computed as the time between consecutive R-peaks in milliseconds. If a participant had IBIs outside of 3 standard deviations (SD) distance from the participant-level average, the trials that included those intervals were excluded from further analysis (1, 1, and 10% of trials were rejected in 3 participants, no trials were rejected in other participants at this step).

#### 2.4.2. Respiration data pre-processing

First, outliers from each 1000 ms moving window of the raw respiration data (mean ∓ 3*SD) were linearly interpolated, using the filloutlier() function in MATLAB. The data was then smoothed by a Savitzky-Golay filter that applies smoothing according to a quadratic polynomial which is fit over a window of 1000 data points. Expiration onsets (peak points on the sinusoidal data) were first automatically determined by the “CARE-rCortex” toolbox in MATLAB (Grosselin et al., 2018), then visually investigated to correct the false positives and negatives. The interval from one expiration onset to the next expiration onset was considered as one respiratory cycle with expiration onset assigned as 0°. If a participant had respiratory cycles with more than two times the median cycle duration, those cycles were not considered for further analysis (mean ± SD = 24 ± 19.7 % of the trials were rejected at this step). 23 out of 27 participants had good quality data used for further analyses (respiratory peaks could not be detected in 4 participants).

#### 2.4.3. EEG data pre-processing

EEG data was pre-processed by using the Fieldtrip software (Oostenveld et al., 2011), EEGLAB toolbox (Delorme & Makeig, 2004), and custom MATLAB scripts. Raw EEG data from each participant was first high-pass filtered at 0.1 Hz, and low-pass filtered at 40 Hz with the basic Finite Impulse Response (FIR) filter of the EEGLAB. Line noise at the 50 Hz and the harmonic frequency of 100 Hz was removed by the CleanLine plug-in of the EEGLAB. Data was then downsampled to 500 Hz, and epoched between -3 and +1.5 seconds with respect to the target stimulus onset. Trials with visible artifacts (i.e., low quality data and blinks in the time window of -1 to +1 seconds with respect to the stimulus onset) were excluded from further preprocessing and analysis (mean ± SD = 8 ± 5% of the trials were rejected at this step).

For each participant, ICA was applied on the remaining epochs. Prior to the ICA, channels with deviant power spectra curve (visibly lower values throughout the power spectra, or excessive high-frequency noise) were also excluded (a minimum of zero and a maximum of two channels were excluded for each participant at this step. ICA was performed by the runica() function on the EEGLAB plug-in. 35 of the ICA components that explain the highest amount of variance in the data were visualized, and the ICLabel component classifier (Pion-Tonachini et al., 2019) was utilized to determine the putative sources of each component. Components with a 90% probability of being a blink or muscle artifact according to the classifier were removed. ICA components were also manually investigated to verify the classification results, and the exclusion criteria were slightly adjusted if necessary (mean ± SD = 9 ± 4.08 ICA components were removed). If there were excluded channels from the ICA, they were spherically interpolated by the pop_interp() function of the EEGLAB plugin at this step.

A two-step baseline correction was applied before obtaining trial-averaged visual evoked potentials (VEPs). First, trials were epoched between -1 and +1 seconds with respect to the target stimulus onset. The timing of the target stimulus onset was shuffled between trials 100 times, and each iteration yielded a surrogate VEP. The average of these surrogates was subtracted from the non-corrected VEP. Surrogate correction was deemed relevant for our paradigm because of a pre-stimulus negative slope in EEG responses (Contingent Negative Variation) that is commonly observed in paradigms where a target stimulus is “contingent” upon a previous warning stimulus (Azzalini et al., 2019; Hillyard, 1969; Steinfath et al., 2024). Finally, the baseline between -100 ms and 0 ms with respect to the target onset was subtracted to obtain the corrected VEP for each participant.

### 2.5. Statistical Methods and Analyses

#### 2.5.1. Respiratory Sinus Arrhythmia (RSA) Estimations

RSA was modeled at the level of each participant by fitting sinusoidal functions on the cluster of data points that correspond to the respiratory phase in radians (0-2π, on the x-axis) and interbeat interval difference (ΔIBI (ms), on the y-axis) at the time of each R-peak (**Fig. 1C**). For the resting state RSA modeling, all R-peaks during the 10-minute baseline recording were used as input. For the RSA during the stimulus anticipation period of the task, the last 3 R-peaks that precede stimulus onset were used separately to track RSA amplitudes and vertical offsets. These non-overlapping data points yielded a beat-by-beat, trial-averaged RSA estimations.

ΔIBI values were first z-scored, and values that deviated more than ±3SDs were excluded from further analyses. Sinusoidal model function was defined as follows:

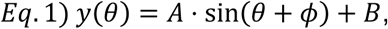

wherein A is the amplitude, Φ is the phase shift, and B is the vertical offset of the sinusoidal curve.

For each set of [respiratory phase, ΔIBI_i_] values used in the RSA estimation, the following loss function L was minimized by using fminsearch() function in MATLAB with the initial guesses of max(ΔIBI)-min(ΔIBI), 0, and mean(ΔIBI) for the parameters A, Φ, and B, respectively:

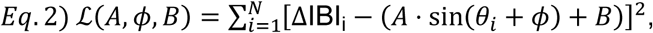

The values of A and B parameters that minimize the loss function were accepted as the estimations for RSA amplitude and respiratory phase-independent cardiac deceleration, respectively. The parameter Φ is also optimized to reach the best possible fit, since the influence of respiratory phase on the heart rate occurs with some delay, which itself may vary between participants (Ben-Tal et al., 2012; Ghibaudo et al., 2023).

The RSA amplitude estimations derived from sinusoidal modeling approach was compared against the Porges-Bohrer method, an established way of calculating RSA amplitude in a time-window of interest. Since the Porges-Bohrer method requires IBI time-series of at least 15-30 seconds, we performed the comparison on the 10-minute resting state data. Details for the implementation of Porges-Bohrer method and its variants are explained elsewhere (Lewis et al., 2012; Porges & Bohrer, 1990). Briefly, the discrete IBI values were interpolated at 4 Hz, then bandpass filtered between 0.12-0.40 Hz. Data was then segmented into 60-second windows with 50% overlap. Variance of each segment was computed and log-transformed. The mean log-transformed variance corresponded to the estimated RSA amplitude of the participant at rest.

#### 2.5.2. Circular Statistics

Uniformity of respiratory phase distributions at time points of interest were tested by using the circular statistics toolbox in MATLAB (Berens, 2009). Respiratory phase of each participant at a given time in the trial (e.g., at the time of the last R-peak before target stimulus onset) was taken as a unit vector, and the vectoral sum from all trials was divided with the number of trials to obtain the participant-level respiratory phase vector. Group-level mean angles and vector lengths were computed by summing and averaging the participant-level vectors. Circular Rayleigh test was utilized to test the null hypothesis of uniform distribution of participant-level vectors in the circular space (Berens, 2009).

#### 2.5.3. Linear Correlations

For the analyses regarding across-participant linear relationships between two variables, the correlation coefficients were computed by fitting a robust linear regression model with the function fitlm() in MATLAB, including an intercept term. Huber’s loss function was used in order to downweigh the effect of data points according to how close they are to being outliers.

Across-participant linear correlations were also computed between time series of participant-level visual evoked potentials from the EEG signal, and the following cardiorespiratory parameters: anticipatory cardiac deceleration, RSA amplitude estimates, and the vertical offset parameter. A cluster-based non-parametric correction method (Maris & Oostenveld, 2007) was applied to determine time windows of the visual evoked potentials that significantly correlated with the aforementioned cardiorespiratory parameters, with critical alpha level set to 0.01.

#### 2.5.4. Statistics on Visual Evoked Potentials

To compare between time series of visual evoked potentials in different trial categories (in trials with visual awareness and no visual awareness of the target, i.e., Hit vs. Miss trials), we used the EEGLAB toolbox (Delorme & Makeig, 2004) and the Fieldtrip software (Oostenveld et al., 2011) in MATLAB to utilize their non-parametric cluster-based permutation tests. Specifically, we used Montecarlo statistics with 1000 iterations for shuffling the data labels, followed by the cluster correction method with critical alpha value of 0.01.

#### 2.5.5. Other Statistical Tests

In order to compare a given cardiorespiratory parameter’s value over the course of stimulus anticipation, repeated-measures ANOVAs were used (i.e., ranova() in MATLAB). If the Mauchly’s test was violated, Greenhouse-Geisser correction was applied. In these cases, the reported p-values correspond to the values after the correction step. When there were two factors in the repeated measures analyses (e.g., two within-participant factors: trial category as Hit vs. Miss, and time), fitrm() function was utilized from MATLAB to fit a repeated measures model. The model was then submitted to the ranova() function for the analysis of variance. Multiple regression analysis was performed with the regress() function in MATLAB.

To compare correlation coefficients obtained separately from different trial conditions, we used the web interface for the cocor package (version 1.1-4) from R (Diedenhofen & Musch, 2015).

## 3. RESULTS

### 3.1. Behavioral Results

Participants detected the threshold-level visual stimulus on average 58.39 ± 2.64% (mean ± SD) of the trials, with an average decision confidence of 69.44 ± 13.96% (mean ± SD). False alarm rate was not higher than 1 per 9 stimulus-absent trials in each experimental block, (see **section 2.1**).

### 3.2. The Role of Respiratory Sinus Arrhythmia (RSA) and Respiratory Phase Alignment in Anticipatory Cardiac Deceleration (ACD)

During the time window that spanned the last three heartbeats before the target stimulus onset (i.e., anticipatory window), a prominent heart rate deceleration was observed (**Fig. 2A -top**). While the positive ΔIBI (IBI_next_ – IBI_previous_) at each of the anticipatory heartbeats indicated the presence of ACD throughout the anticipatory window, the amount of ACD also increased over the course of the three heartbeats (one-way repeated-measures ANOVA; F(2,22)=18.38, p=6.33x10^-5^).

During the same time window, we found that participants displayed significant respiratory phase alignment at the time points of each pre-stimulus heartbeat (**Fig. 2A -bottom**). **Supplementary Table 1** shows the group-level respiratory phase vector lengths and angles, as well as the results from the Rayleigh’s test for uniform distribution of the participant-level average vectors at the group-level. As can be seen both qualitatively from **Fig. 2A** and quantitatively from **Supplementary Table 1**, participants aligned their respiration to progressing phases of exhalation during the anticipation period.

As expected, analysis of the resting-state data showed that the Respiratory Sinus Arrhythmia (RSA) effect was prominent at the group-level, with a net cardiac deceleration (positive ΔIBI values) during exhalation, and a net cardiac acceleration during inhalation (**Fig. 2B**).

To investigate the potential contribution of RSA on the cardiac deceleration during the stimulus anticipation window, we modeled the RSA at resting-state for each participant by fitting a sinusoidal curve on the data points corresponding to the heart rate change around the R-peaks of interest and the respiratory phase at these time points (see **Fig. 1C** and **section 2.5.1** for more details). RSA amplitudes were first estimated from the resting-state data with our sinusoidal models for each participant (**Fig. 2C**, see **Supplementary Table 2** for participant-level model outputs), and compared to the traditional Porges-Bohrer RSA amplitude estimation method (**Fig. 2D**) to verify the accuracy of our sinusoidal models. RSA estimations derived from the two methods correlated significantly (r(22) = 0.83, p = 1.84x10^-5^; **Fig. 2D**).

Sinusoidal modeling was then performed on data points corresponding to the R-peaks from the anticipatory window from the task trials (specifically the last three R-peaks before stimulus onset in each trial, see **Fig. 1C**), and the resulting RSA estimates were compared against the ACD amplitudes seen in the same window (**Fig. 2E**, see **Supplementary Table 3** for participant-level model outputs)). RSA amplitudes in the stimulus anticipation window correlated significantly with the amplitude of the ACD across participants (r(22) = 0.65, p = 0.09x10^-2^; **Fig. 2E**).

A multiple regression model was also run with the baseline and anticipatory RSA values from each participant as the two predictor variables, and the total ACD as the response variable. Results showed a significant correlation between the total ACD and the predictor RSA variables (r(22) = 0.68, p = 0.07x10^-1^). In addition, the baseline and anticipatory RSA amplitudes of the participants correlated significantly (r(22) = 0.87, p = 7.14x10^-8^), showing that the participants with a higher RSA amplitude at rest also displayed a higher RSA amplitude during stimulus anticipation. Combined with the previous results of respiratory phase alignment to exhalation (see **Fig. 2A-B**), these results suggest that the RSA amplitude contributes to the cardiac deceleration response during stimulus anticipation, raising the question of whether ACD can be fully explained by the respiratory phase-dependent effect of RSA.

**Figure 2:**
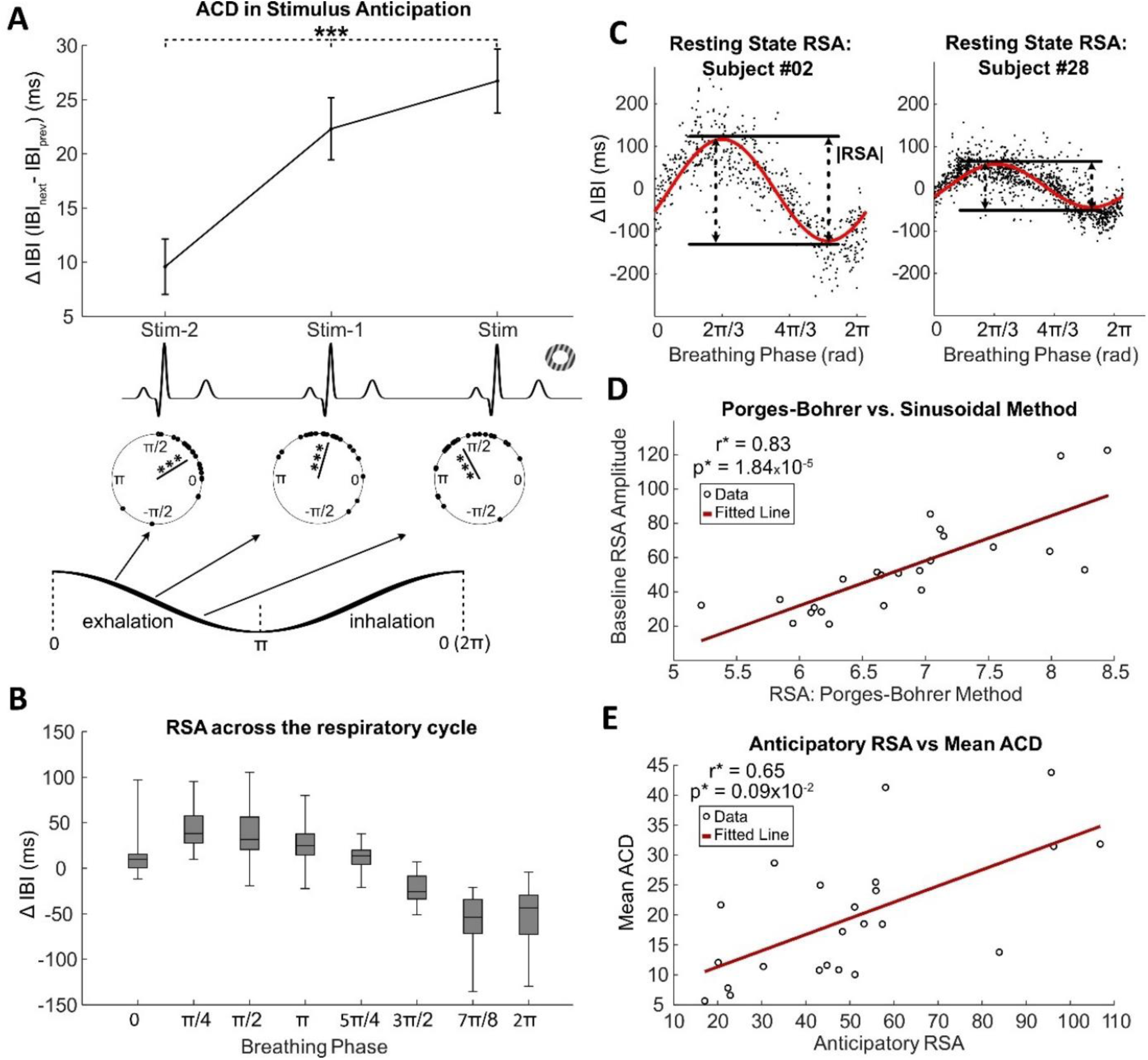
The role of RSA in ACD. **A)** Anticipatory Cardiac Deceleration (ACD) during stimulus anticipation (top), and group-level respiratory phase-locking at the time of each pre-stimulus heartbeat (bottom). Each point on the circular plots represents the mean respiratory phase direction of a participant at the corresponding time point. (Error bars correspond to s.e.m., ***: p<0.001 on all sub-figures; the doughnut-shaped grating represents the stimulus onset which comes after the R-peak named “stim”). **B)** Illustrative plot for the effect of RSA on heart rate at the group-level. The plot was constructed from resting-state R-peaks (bars indicate minimum and maximum data values, boxes correspond to the interquartile range, and the median value is denoted with a black horizontal line within each box). **C)** Single-participant examples of resting-state RSA curves. Sinusoidal functions were fitted to the cluster of data points corresponding to each R-peak during the 10-minute resting-state recordings (|RSA|: RSA amplitude estimate). **D)** Comparison of RSA estimation methods by linear correlation. Our sinusoidal model is compared to the established Porges-Bohrer method. **E)** Correlation between the anticipatory RSA and task-related ACD at the across-participant level (*: r (correlation coefficient) and p-values correspond to robust linear regression, **see section 2.5.3**).

### 3.3. Respiratory Phase-Dependent and Phase-Independent Contributors to Anticipatory Cardiac Deceleration (ACD)

Next, we investigated whether ACD could be fully explained by respiratory phase-alignment to exhalation and the resulting cardiac deceleration due to Respiratory Sinus Arrhythmia (RSA). We first categorized the trials according to whether the R-peaks in the stimulus anticipation window (“stim-2”, “stim-1”, and “stim”) occurred during exhalation or inhalation. As expected, we observed an increasing amount of average cardiac deceleration from the beginning to the end of the anticipatory window in exhalation (**Fig. 3A**). In the trials with anticipatory R-peaks in inhalation, we observed a net cardiac acceleration; but the amount of cardiac acceleration decreased and was almost cancelled out at the last R-peak before the target stimulus onset (“stim” R-peak, denoted by inh-0 in **Fig. 3A**). During the resting-state, participants’ cardiac acceleration during inhalation was 30.30±15.3 ms (mean ± SD), whereas the anticipatory cardiac acceleration during inhalation (the average cardiac acceleration around R-peaks denoted by “inh-2”, “inh-1”, and “inh-0” in **Fig. 3A**) was limited to 12.69±13.48 ms (mean ± SD). A Wilcoxon signed-rank test showed that the RSA-based cardiac acceleration during inhalation was significantly attenuated during stimulus anticipation (n = 23, z = -4.11, p = 4.03x10^-5^), suggesting the presence of a respiratory phase-independent factor that decelerates the heart irrespective of the respiratory phase.

Next, we sought to quantify the respiratory phase-independent contribution to the total ACD systematically. While the amplitude of the sinusoidal RSA functions (**Fig. 3B – top**) predicted the RSA amplitude accurately (see **Fig. 2**), we postulated that the vertical offset (variable “B”) of these curves could be used to predict the baseline tendency of the heart rate to decrease or increase, irrespective of the respiratory phase (**Fig. 3B -bottom**; positive and negative vertical offset would be interpreted as a baseline decrease and increase in heart rate, respectively).

In addition to the significant correlation between the baseline and anticipatory RSA, we observed that the RSA amplitudes in the two periods did not differ significantly (**Fig. 3C** - **left**; one-way repeated-measures ANOVA; F(3,22)=2.40, p=0.11). On the other hand, the vertical offset parameter (“B”) was not different from zero during the resting-state, as expected in the absence of any temporary heart rate modulations due to anticipation (Wilcoxon signed-rank test, n=23, z = -1.19, p = 0.24). Unlike in the resting-state; B was visibly above zero during stimulus anticipation, and it increased significantly within the anticipatory window (**Fig. 3C** - **right**; one-way repeated-measures ANOVA; F(3,22)=27.94, p=9.41x10^-9^).

Since both the RSA amplitude and the vertical offset (variable B) are measured on the vertical axis, we controlled if they co-fluctuate across different states and time points. Therefore, we first tested beat-by-beat co-fluctuations of the two variables from the “stim-2” to “stim-1” R-peak (**Fig. 3D – top**; r(22) = 0.29, p = 0.21), and from the “stim-1” to “stim” R-peak (**Fig. 3D – bottom**; r(22) = -0.10, p = 0.75). In addition, the average RSA amplitude during stimulus anticipation did not correlate with the upward shift in the vertical offset (ΔB) during the last three heartbeats before stimulus onset (**Fig. 3E**; r(22) = -0.02, p = 0.59). The results show that the two variables did not covary, hinting towards at least partially independent underlying mechanisms.

**Figure 3:**
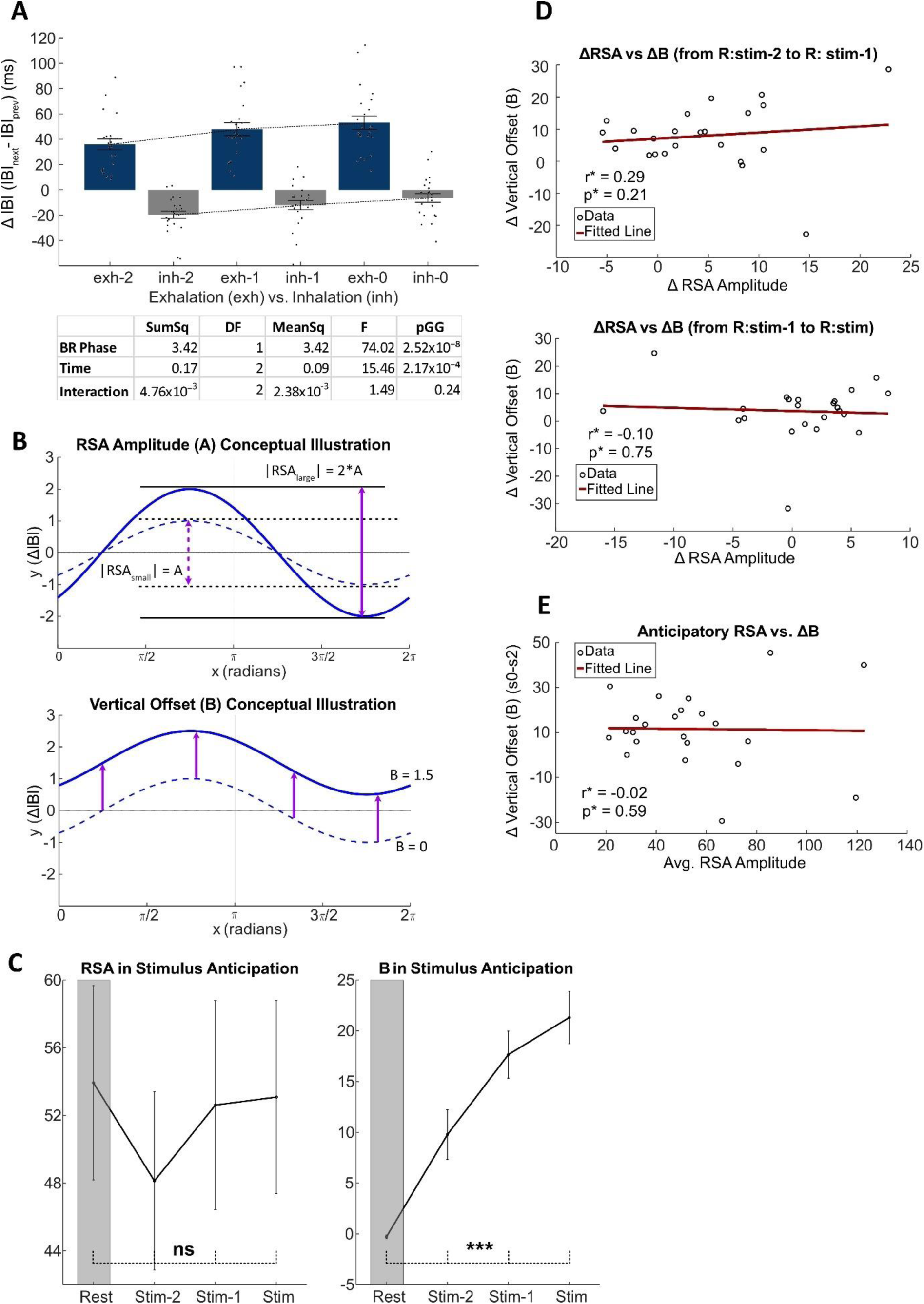
Quantifying respiratory phase-dependent and phase-independent heart rate modulations during stimulus anticipation. **A)** ACD in stimulus anticipation during exhalation and inhalation. Numbers on the x-axis labels represent the timing of the R-peak with respect to the stimulus onset (Error bars indicate s.e.m.). “exh-0” and “inh-0” correspond to “R:stim”, the last R-peak before stimulus onset. Rest of the annotations follow accordingly (see Fig. 1B). Table corresponds to the results from two-way repeated measures ANOVA test (Within-subject dependent factors are Respiratory Phase (“BR Phase”, and Time); SumSq: sum of squares, MeanSq: mean square, pGG: p-value after Greenhouse-Geisser correction). **B)** Modeling the effects of respiratory phase-dependent and phase-independent contributions to the ACD. The amplitude (parameter A) and vertical offset (parameter B) of the fitted sinusoidal RSA curves are postulated as the respiratory phase-dependent and phase-independent components of ACD, respectively. The plots are solely for illustrative purposes. **C)** Parameters A (i.e., RSA; left) and B (right) during resting-state and stimulus anticipation. Progress of the parameter values starting from the resting state (Rest) and throughout the stimulus anticipation phase (from “R:Stim-2” to “R:Stim”; ***: p<0.001, ns: p>0.05, bars indicate s.e.m.). **D)** Statistically independent modulations of the parameters A and B throughout stimulus anticipation. From “R:stim-2” to “R:stim-1” (top), from “R:stim-1” to “R:stim” (bottom). (*: r and p-values are derived from robust linear regression, see **section 2.5.3**). **E)** Across-participant independence of average RSA amplitude in stimulus anticipation and ΔIBI from “R:stim-2” to “R:stim”.

On the other hand, the RSA amplitudes during both resting-state and stimulus anticipation were positively correlated with the average vertical offset (B) during stimulus anticipation (**Supplementary Fig. 1**; r(22) = 0.64, p = 0.01x10^-1^ for resting-state RSA; r(22) = 0.59, p = 0.04x10^-1^ for anticipatory RSA). However; the beat-by-beat increase in the vertical offset (ΔB) could not be explained by the respiratory phase-dependent effect of RSA as explained above (see **Fig. 3D-E**).

### 3.4. Differential Contributions of Respiratory Phase-Dependent and Phase-Independent Heart Rate Modulations to Visual Stimulus Detection

In our perceptual task, the participants were asked to report whether they detected the threshold-level target visual stimulus at the end of each trial. We next investigated how the prominent cardiac deceleration during target anticipation (ACD), or specifically the respiratory phase-dependent (RSA) and phase-independent (vertical offset, or parameter “B”) factors contributing to the total ACD were related to visual stimulus detection. Below, we name the trials in which the participants correctly detected the target as “Hit”, and the ones in which they did not see the presented target as “Miss”.

First, we observed that the total ACD during stimulus anticipation did not differ between Hit and Miss trials (F(1,22) = 0.56, p = 0.46, **Fig. 4A**). On the other hand, the RSA amplitude was lower in Hit trials than in Miss trials throughout the stimulus anticipation period, reflected by the main effect of “Detection” in the corresponding two-way repeated measures ANOVA (within-subject factors: “Detection” and “Time”; F(1,20) = 5.16, p = 0.03, **Fig. 4B**).

We have shown that the vertical offset component (B) increased significantly within the stimulus anticipation period (see **Fig. 3C -right**). Here, we compared this upward shift in Hit and Miss trials, and observed that the vertical offset shifted more in Hit trials than in Miss trials to a small, but statistically significant extent (Wilcoxon signed-rank test; n = 23, z = 2.07, p = 0.04, CI for the difference between categories = [0.21 6.38]; **Fig. 4C and 4D -left**). The average vertical offset during stimulus anticipation did not differ between Hit and Miss trials (Wilcoxon signed-rank test; n = 23, z = -0.36, p = 0.72, CI for the difference between categories = [-0.99 0.93]; **Fig. 4D -right**), suggesting that a beat-by-beat increase in vertical offset –rather than its average value– during stimulus anticipation is specifically related to perceptual performance. Overall, these results suggest a decrease in the respiratory phase-dependent RSA amplitude and a steeper increase in phase-independent heart rate deceleration during the anticipation phase in Hit vs. Miss trials.

We also median-splitted the confidence ratings from each participant to create trial conditions of low- and high-confidence, and compared the same set of cardiorespiratory parameters between these two conditions. However, neither the total ACD, nor the RSA amplitude, nor ΔB differed in amplitude between low- and high-confidence conditions (**Supplementary Fig. 2**).

Finally, we investigated the correlations between the cardiac deceleration parameters (total ACD, parameters A and B) and Visual Awareness Negativity (VAN), a visual evoked potential that is observable at occipital EEG electrodes or MEG sensors and is associated with the conscious perception of –and/or attention to– visual stimuli (Bola & Doradzińska, 2021; Li et al., 2014; Mazzi et al., 2020).

**Figure 4:**
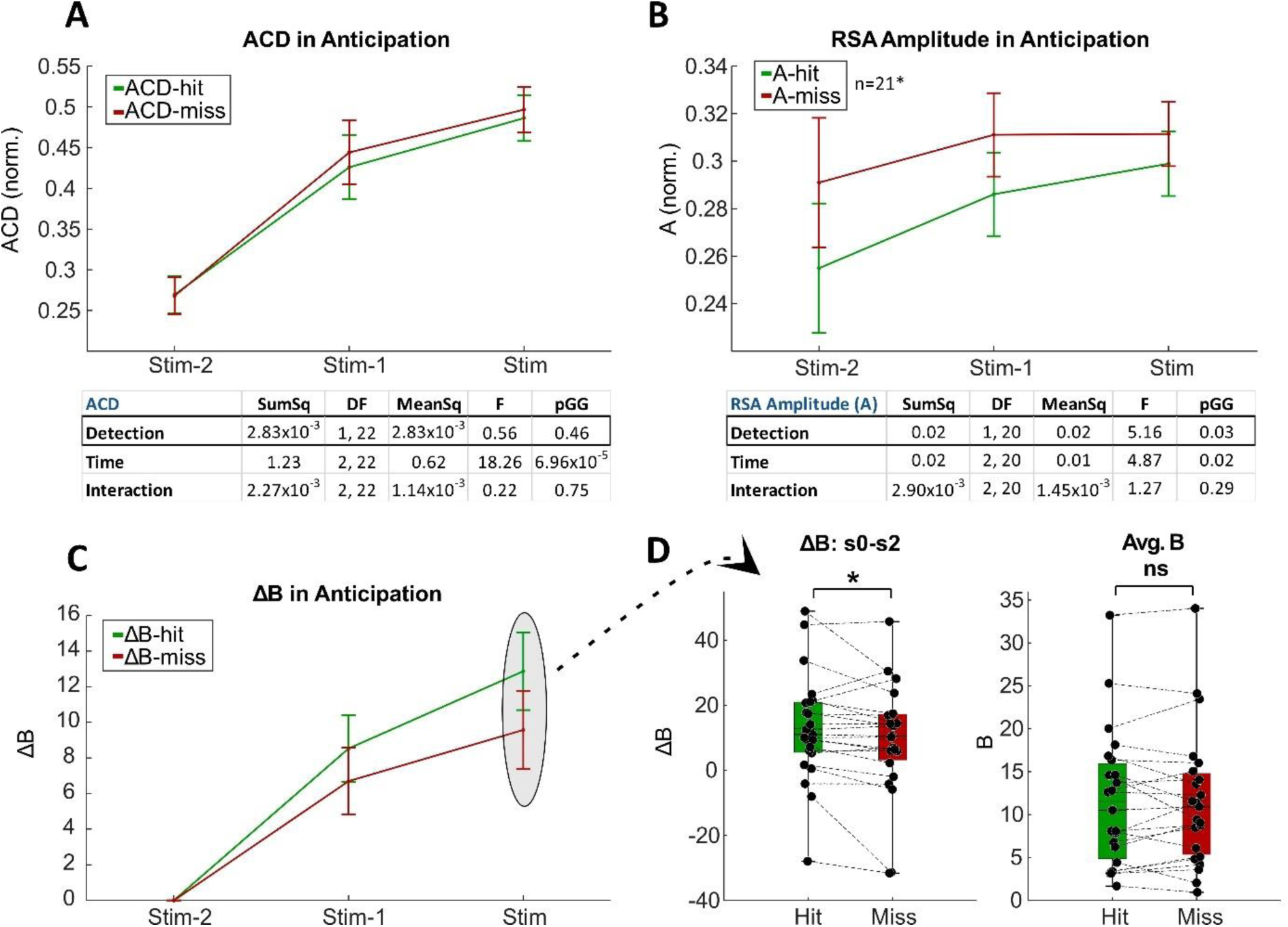
RSA and respiratory phase-independent cardiac deceleration in Hit and Miss trials. **A)** Total cardiac deceleration, **B)** Estimated RSA amplitude during stimulus anticipation in Hit (successful detection) and Miss (false rejection) trials. Tables in A and B show the results of the two-way repeated measures ANOVA analysis on the effects of Stimulus Detection (Hit vs. Miss) and Time on the total ACD and estimated RSA amplitudes, respectively. (*: two participants had to be excluded due to outlier values; SumSq: sum of squares, MeanSq: mean square, pGG: p-value after Greenhouse-Geisser correction). **C)** Progress of the vertical offset parameter within stimulus anticipation. The time point of “R:stim-2” heartbeat was taken as reference. Error bars in **A**,**B**, and **C** correspond to within-participant 95% confidence intervals (Cousineau, 2005; Morey, 2008). **D)** Comparison of shifts in vertical offset (ΔB) from the beginning to the end of stimulus anticipation in Hit and Miss trials (left), and the average vertical offsets during stimulus anticipation (right). (Wilcoxon signed-rank test, *: p<0.05, ns: p>0.05; error bars indicate the range of data points).

Since the VAN is associated with conscious perception –thus expected to be significantly attenuated in Miss trials– we first investigated the time window in which the evoked potential on the occipital electrodes (the average activity from the O1, Oz, and O2 electrodes) differed significantly between Hit and Miss trials. After cluster-based correction, this time window corresponded to 188-318 ms after the target onset (**Fig. 5A**). The linear correlation analysis across-participants showed that the average VAN amplitude in this time window was significantly correlated with the total cardiac deceleration in the anticipation window (ACD) (r(22) = 0.61, p = 0.02x10^-1^; **Fig. 5B**). The same correlation was significant in Hit trials (r(22) = 0.52, p = 0.01; **Fig. 5C-left**), but not in Miss trials (r(22) =0.25, p = 0.25; **Fig. 5C-right**). A direct comparison of correlation coefficients revealed that the correlation coefficient in Hit trials was indeed significantly more negative than in Miss trials (comparison of nonoverlapping and dependent correlation coefficients (Diedenhofen & Musch, 2015); Dunn and Clark’s z(22) = -1.86, p = 0.03, one-tailed).

To control for the specificity of the time window in which the visual evoked potentials correlated with ACD across-participants, we investigated all time points of the evoked potential responses from the occipital electrodes (i.e., blind to the time window of the VAN) and determined the time points of evoked responses that showed a significant correlation with the cardiac deceleration amplitude after temporal cluster-correction. We found that the only time window with a significant correlation between the visual evoked potential and ACD was 244-302 ms after stimulus onset (**Fig. 5D**), which is completely within the time window in which the VAN was observed (188-318 ms after stimulus onset, see **Fig. 5A**).

Since visual evoked potentials can also reflect the contrast of the stimulus, we ensured that the stimulus contrasts did not differ between trial categories: we downsampled the Hit trials of each participant by sorting from the lowest to highest stimulus contrast, and using the first ones until the trial number for the Miss trials was reached. The stimulus contrast did not differ between the downsampled Hit and Miss trials (**Supplementary Fig. 3A**), and the VAN amplitudes did not correlate with the average stimulus contrasts shown to the participants (**Supplementary Fig. 3B**).

The gradual increase in the respiratory phase-independent heart rate modulator (ΔB) also showed a significant correlation with the visual evoked response in the time window corresponding to the VAN (r(22) = -0.41, p = 0.04; **Fig. 5E – left**), and this significance was present in Hit trials (r(22) = -0.44, p = 0.03; **Fig. 5E – middle**) but not in Miss trials (r(22) = - 0.28, p = 0.39; **Fig. 5E – right**). However, the direct comparison of the correlation coefficients from the Hit and Miss trials did not reveal a significant difference, in contrast to the total ACD (one-tailed comparison of nonoverlapping, and dependent correlation coefficients; Dunn and Clark’s z(22) = -1.19, p = 0.12). On the other hand, VAN amplitudes did not significantly correlate with the RSA amplitudes (**Fig. 5F**), suggesting that the effect of ACD on VAN is likely mediated by the respiratory phase-independent contributor rather than RSA.

**Figure 5:**
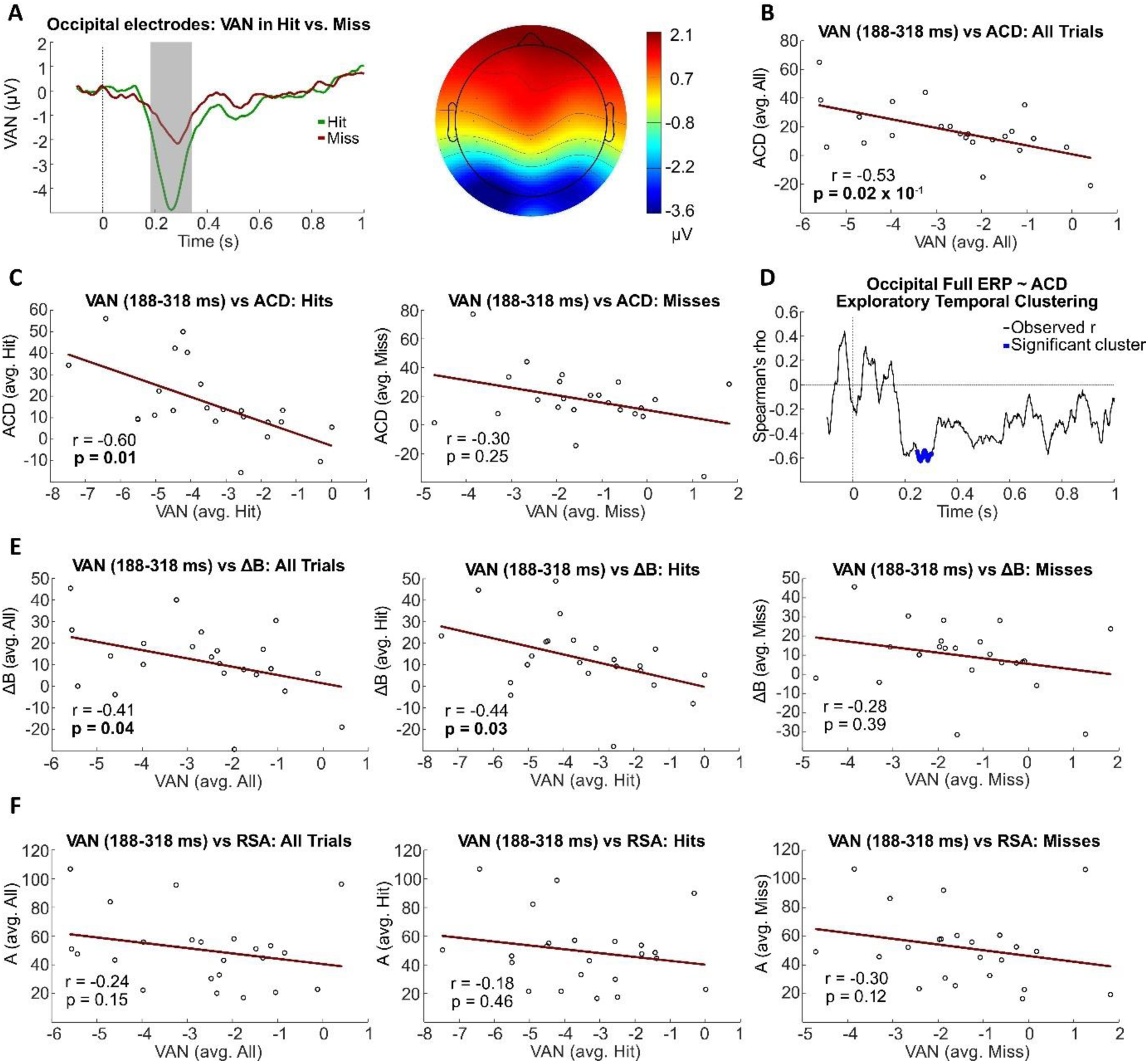
Across-participant correlations between cardiac deceleration parameters and Visual Awareness Negativity (VAN). **A) Left:** Average visual evoked potentials at the occipital electrodes (Oz, O1, O2) in response to the target stimulus onset. Grey rectangle denotes the time window of 188-318ms (with respect to stimulus onset, dashed vertical lines) in which the difference between the ERP in Hit and Miss trials was significant. **Right:** Average scalp topography of the significant time window (unit of heatmap is in microvolts). **B)** Across-participant linear correlation between the average VAN amplitude and ACD for each participant. **C)** Same linear correlations as in **B**, but using the values from only Hit (left) and only Miss (right) trials (r: Spearman’s correlation coefficient, p: p-value for the significance of the correlation based on robust linear fitting). **D)** Exploratory analysis of significant time window clusters for the across-participant correlation between the full visual evoked potential time window (0-1s with respect to the stimulus onset) and ACD amplitude in Hit trials. The thick blue line denotes the significant cluster of Spearman’s correlation coefficients between 244-302 ms with respect to the stimulus onset (dashed vertical lines). **E)** Across-participant correlations between VAN amplitudes and the respiratory phase-independent (vertical offset, ΔB) and **F)** phase-dependent (RSA) contributors to the anticipatory heart rate modulation. All p-values correspond to robust linear regression).

## 4. DISCUSSION

Anticipatory Cardiac Deceleration (ACD) is a temporary decrease in heart rate, observed during perceived threat or anticipation of a task-relevant event (Ribeiro & Castelo-Branco, 2019; Skora et al., 2022). Even though heart rate is also constantly modulated as a function of the respiratory phase due to Respiratory Sinus Arrhythmia (RSA; or respiratory Heart Rate Variability (Menuet et al., 2025)), ACD is typically quantified without taking RSA into account. Here, we demonstrated that a novel trial-averaged RSA estimation method can simultaneously estimate the respiratory phase-independent cardiac deceleration.

In our perceptual task, we found a respiratory phase alignment to advancing stages of exhalation during stimulus anticipation. This alignment in itself supports cardiac deceleration via RSA, due to the vagal parasympathetic input being disinhibited during exhalation (Berntson et al., 1993; Elstad et al., 2018). We showed that the RSA amplitude is indeed correlated with the total ACD, suggesting at least a facilitatory role of RSA on ACD, implemented by the respiratory phase alignment to advancing stages of exhalation during stimulus anticipation.

However, ACD is observed across many different experimental paradigms and settings (Alam et al., 2023; Motyka et al., 2019; Park et al., 2014; Skora et al., 2022) which do not necessarily observe the same respiratory phase alignment that supports cardiac deceleration. Thus, we hypothesized that the respiratory phase-dependent effect of RSA should not be the sole contributor to the ACD. In line with this, our results showed that RSA-based cardiac acceleration during inhalation was also gradually attenuated during stimulus anticipation, suggesting that the cardiac deceleration was not solely due to the exhalation-specific effect of RSA. The positive and increasing vertical offset in the sinusoidal RSA functions during stimulus anticipation also supported the presence of respiratory phase-independent cardiac deceleration. Importantly, we were able to quantify this component while taking into account the RSA amplitude of the participants, and found that the RSA amplitude and non-respiratory cardiac deceleration do not covary beat-by-beat during stimulus anticipation. However, the average vertical offset value (B) during stimulus anticipation was positively correlated with the average RSA amplitudes during resting-state and stimulus anticipation. These results suggest the co-existence of two partially independent modulators of heart rate during stimulus anticipation. On the one hand, RSA is typically considered to be a marker of overall cardiac parasympathetic tone (Lane et al., 1992; Thayer & Lane, 2000), and is positively associated with HRV and flexibility of cardiac modulations based on environmental demands (Porges, 1995; Siciliano et al., 2022). Our data supports this view since we observed that the participants with higher baseline RSA amplitudes also displayed a larger average of temporary cardiac deceleration (vertical offset, or B) during stimulus anticipation, which exemplifies a flexible cardiac modulation based on a task-relevant demand. On the other hand, we observed that the vertical offset gradually increased from the beginning towards the end of the anticipatory window. This increase was independent of the RSA amplitude, unlike the average amount of non-respiratory cardiac deceleration during stimulus anticipation. The underlying physiological mechanism of ACD is suggested to be a top-down parasympathetic activation initiated by a set of cortical centers –corresponding to the Central Autonomous Network regions– in response to the task-related attentional demands (Battaglia et al., 2025; Skora et al., 2022; Somsen, Jennings, & Van der Molen, 2004), which likely functions independent of the respiratory vagal input to the heart. Based on our results, the positive vertical offset throughout stimulus anticipation might reflect the respiratory phase-independent cardiac deceleration, the amplitude of which seems to be predictable by the baseline RSA amplitudes of the participant. The gradual increase in the amount of this deceleration during stimulus anticipation might serve to meet the temporal attentional demands which are maximized towards the end of the anticipatory period, and is no longer correlated with the underlying RSA amplitude of the participant.

Furthermore, we compared the RSA amplitude and the vertical offset component during stimulus anticipation between Hit and Miss trials. We found that the non-respiratory cardiac deceleration increased more throughout the anticipation phase, whereas the RSA amplitude was lower in Hit trials than in Miss trials. Previous studies on sensory perception have also observed and analyzed cardiac deceleration during stimulus anticipation (Grund et al., 2022; Park et al., 2014), finding similar pre-stimulus ACD amplitudes in their Hit and Miss trials, consistent with our results. However; those studies did not consider RSA and non-respiratory modulations of the heart rate in Hit vs. Miss trials. Here we show that the two components are modulated in opposite directions during stimulus anticipation in Hit and Miss trials, thus possibly cancelling out each other’s effects on the heart rate. Given that the group-level respiratory phase alignment profiles would likely differ between studies, we suggest that controlling for respiratory phase-dependent effects on the heart might increase the replicability of ACD-related findings across studies. Specifically, if the respiratory phase aligned to inhalation during anticipation, co-existence of RSA-based cardiac acceleration and non-respiratory cardiac deceleration might lead to significant underestimation of the cognitive control of heart rate in the anticipatory window. Therefore, it would be interesting to see similar characterizations of RSA-based and non-respiratory heart rate modulations in different anticipation- or threat-related paradigms. We believe that our simple sinusoidal modeling approach to simultaneously estimate RSA amplitudes and non-respiratory heart rate modulations provides a useful tool for researchers to investigate the heart rate dynamics in any given task context while controlling for respiration-dependent oscillations of the heart rate.

Porges’ two-component model of attention suggests that while sustained attention is marked by a reduction in RSA amplitude, reactive attention leads to temporary cardiac deceleration as part of an orienting reflex towards the target event (Porges, 1992; Somsen, Jennings, & Van Der Molen, 2004). Relatedly, performance of a cognitive task is typically associated with a decrease in RSA, and the amount of decrease in RSA is used as a marker for attention (Somsen, Jennings, & Van der Molen, 2004; Suess et al., 1994). In this study, we did not find a significant reduction in the RSA amplitude during stimulus anticipation with respect to the resting-state, but we found that the anticipatory RSA amplitude was lower in Hit trials than in Miss trials. The relative decrease in RSA in Hit trials supports the association between RSA attenuation and allocated attention in the sense that attentional demands of the task were likely met more efficiently by the participants in these successful trials. We also observed that the non-respiratory heart rate modulation increased gradually during stimulus anticipation, and that this increase was more accentuated in Hit trials. This result supports the idea that the parasympathetic break on the heart may be functioning in a precision weighting mechanism wherein a decreased precision is assigned to interoceptive signals and an increased precision to exteroceptive signals (Roelofs, 2017; Skora et al., 2022), and that this trade-off is implemented more efficiently in trials with successful visual stimulus detection.

The Visual Awareness Negativity (VAN) is a visual stimulus-evoked potential, typically observed at the occipital scalp EEG electrodes or MEG sensors 200-300 ms following stimulus onset. The VAN is hypothesized to be an early marker of visual awareness (Eklund & Wiens, 2018; Mazzi et al., 2020; Bola & Doradzińska, 2021). Here, we found an across-participant correlation between the ACD and VAN amplitudes, specific to Hit trials. Upon looking at the entire time window of the visual evoked potentials until 1 second after the stimulus onset, we saw that this correlation was specific to the time window of VAN. We also verified that the evoked potentials were not correlated with the average stimulus contrasts shown to each participant. We see that a similar correlation is also present between the slope of the non-respiratory cardiac deceleration (ΔB) and VAN amplitudes, but not between RSA and VAN. This supports the aforementioned idea that the non-respiratory heart rate modulation might be more tightly related to a cognitive top-down parasympathetic input on the heart due to stimulus anticipation, while RSA is a task-irrelevant and tonic modulator of the heart rate.

We reported a slightly more pronounced across-participant correlation between the total ACD and the VAN, compared with the correlation between the non-respiratory cardiac deceleration (ΔB) and VAN. This seems to be in contrast with our findings on the behavioral readout of stimulus detection (Hit vs. Miss trials), wherein we reported a specific effect of ΔB, but not of the total ACD. Firstly, it should be noted that VAN is a neural correlate of conscious perception, but a negative potential deflection at the same time window is not completely absent in the trials with low or no awareness of the target stimulus (Eklund & Wiens, 2018; Koivisto & Grassini, 2016). This was also replicated in our study, with a significantly weaker –but not absent– negative deflection around 200 ms after stimulus onset in the Miss-compared to Hit trials. Given the similarity in the shape of the VAN between Hit and Miss trials, the relationship between cardiac deceleration parameters and the VAN is not expected to be directly translated to the binary parameter of stimulus detection performance, or vice versa. It is also important to note that the correlation between the ΔB and VAN was significant only in the Hit trials as well, even though the direct correlation coefficient comparison between Hit and Miss trials remained as a non-significant trend. We also report that the across-participant correlations between the cardiac deceleration parameters and VAN are statistically significant but relatively small effects. Thus, future studies should test the generalizability of the relationship between cardiorespiratory parameters during stimulus anticipation and stimulus evoked potentials in the cortex, especially under different experimental paradigms and respiratory phase-alignment profiles.

A few other points on the generalizability of our results across different experimental paradigms should be noted. Firstly, ACD has been widely studied in motor tasks with high response time pressure, and the amplitude of ACD is consistently associated with faster response timing or higher motor accuracy (Alam et al., 2023; Ribeiro & Castelo-Branco, 2019; Skora et al., 2022; Somsen, Jennings, & Van Der Molen, 2004). Thus, ACD is postulated to be a marker of action preparation (Hashemi et al., 2019; Skora et al., 2022). Here, we instead utilized a perceptual task with a low response time pressure (i.e., the response time was not relevant to the task). We still observed ACD prominently, and found that even though the total amount of ACD was not directly associated with better perceptual performance, the respiratory phase-dependent and phase-independent contributors to the total ACD were associated with visual detection in opposite directions. Future studies could investigate similar dissociations between respiratory and non-respiratory cardiac deceleration during action preparation, and further elucidate the role of ACD under different task requirements. It should also be noted that even though fast responses are not relevant to performance in our task, preparation for the button presses could still have partially contributed to the total ACD that we observed. Our findings on the relevance of cardiac deceleration parameters to both perceptual performance and visual awareness negativity suggest that ACD does not only serve motor planning and action preparation. However, it would be interesting to see how cardiac deceleration, and specifically respiratory phase-dependent and phase-independent contributors, are related to perceptual performance in a paradigm with a significantly delayed (at least a couple of seconds instead of the 0.3-0.7s in our task) or completely absent motor report.

Secondly, we used a sequence of three heartbeats before stimulus onset for our ACD analyses because this was the maximum number of R-peaks in the pre-stimulus window for most participants. Since cardiac deceleration may be confined to earlier or later stages of longer anticipatory windows (Somsen, Jennings, & Van der Molen, 2004), interactions between RSA and non-respiratory cardiac decelerations could also be compared between anticipatory periods of different lengths. Relatedly, our pre-stimulus window included a total of 0.9-second jitter (i.e., temporal uncertainty; 0.7 seconds in the “Fixation” and 0.2 seconds in the “Warning” period). Previous research suggests that anticipatory windows of fixed length across trials might lead the cardiac deceleration to be steeper and limited to the very end of the anticipatory window (Jennings et al., 1987; Somsen, Jennings, & Van der Molen, 2004), whereas anticipatory windows of variable length yielded more irregular patterns of cardiac deceleration over the course of anticipation (Somsen, Jennings, & Van der Molen, 2004; Van Der Molen et al., 1987). Therefore, the functional relevance of cardiac deceleration parameters (both RSA and non-respiratory factors) in different time windows during stimulus anticipation under various lengths and temporal uncertainties remain to be investigated.

### 4.1. Conclusions

Here we presented a simple sinusoidal modeling approach to estimate trial-averaged Respiratory Sinus Arrhythmia (RSA) amplitudes in various time points of interest from completely non-overlapping data points, while simultaneously extracting non-respiratory modulations of the heart rate. We dissociated the two contributors to the overall Anticipatory Cardiac Deceleration (ACD) in a visual threshold-level stimulus detection task and found that decreased RSA and steeper non-respiratory cardiac deceleration are associated with successful detection. In the EEG, we further found larger Visual Awareness Negativity (VAN) induced by detected stimuli in the participants with a higher ACD. In conclusion, our results support a contribution of cardiac deceleration (especially the non-respiratory component) to visual perception, and highlight the importance of taking respiration into account while investigating heart rate modulations. The role of respiratory and non-respiratory cardiac deceleration in anticipation of different target events (e.g., motor actions or cognitive tasks with varying difficulties) and different anticipation windows remain to be elucidated, and we suggest that our sinusoidal modeling approach provides a useful tool to investigate these questions further.

## Authors contributions

Conceptualization: EK, CMS, MW. Data acquisition: EK, SC. Data curation: EK, SC. Software: EK, SC. Formal analysis: EK, SC. Funding acquisition: CMS, MW. Project Administration: CMS, MW. Supervision: CMS, MW. Visualization: EK, SC. Writing—Original draft preparation: EK. Writing—Review & editing: EK, CMS, MW.

## Acknowledgements

We thank Carsten Schmidt-Samoa for providing scripts on the online control of gaze fixation, Aishwarya Bhonsle and Shamim Sasani Ghamsari for helpful discussions, and Severin Heumüller and Daniela Proto Salvador for providing IT support.

## Declaration of generative AI and AI-assisted technologies in the writing process

During the preparation of this work, we used ChatGPT 4 in order to facilitate some aspects of data analysis implementations in MATLAB. After using this tool/service, we reviewed and edited the code and take full responsibility for the content of the published article.

## Data and code availability statement

The datasets generated and analyzed for the current study, and the corresponding code, are available from the corresponding author on request. Code for running the visual perception task, physiological data pre-processing and analysis can be found in this public GitHub repository: https://github.com/ege-kingir/VisualThresholdDetection

