## Supplementary Information for "Disentangling *Respiratory Phase-Dependent* and *Anticipatory* Cardiac Deceleration in a Visual Perception Task"

**Supplementary information contains:**

3 Supplementary Tables

3 Supplementary Figures

**Supplementary Table 1:** Group-level breathing phase vector angles, lengths, and the results from the Rayleigh's test against uniform distribution (rad: radians, deg: degrees).

| Heartbeat (time point) | Vector Angle (rad, deg) | Vector Length | Rayleigh's z-value | p-value |
|---|---|---|---|---|
| Stim-2 | (0.55, 31.75) | 0.80 | 14.80 | $1.69x10^{-8}$ |
| Stim-1 | (1.29, 74.05) | 0.83 | 15.88 | $2.90x10^{-9}$ |
| Stim | (2.06, 118.47) | 0.77 | 13.51 | $1.21x10^{-7}$ |

**Supplementary Table 2:** Parameters corresponding to the best sinusoidal model for each participant's 'breathing phase ~ interbeat interval' curve based on the R-peaks from the 10-minute *baseline* recording. (A: amplitude, B: vertical offset, φ= phase shift, $R^2$: R-squared (coefficient of determination), $SS_{res}$: residual sum of squares, $SS_{tot}$: total sum of squares)

| **Subj. ID** | **A** | **B** | **ϕ** | **$R^2$** | **$SS_{res}$** | **$SS_{tot}$** |
|---|---|---|---|---|---|---|
| 1 | 32.18 | 0.27 | -0.34 | 0.63 | $1.86x10^5$ | $4.98x10^5$ |
| 2 | 119.49 | -2.92 | -0.43 | 0.72 | $1.53x10^6$ | $5.50x10^6$ |
| 3 | 50.89 | 0.54 | -0.47 | 0.71 | $3.45x10^5$ | $1.18x10^6$ |
| 4 | 66.18 | 0.02 | -0.60 | 0.70 | $6.50x10^5$ | $2.15x10^6$ |
| 5 | 27.89 | 0.04 | -0.78 | 0.56 | $1.89x10^5$ | $4.33x10^5$ |
| 6 | 85.47 | 0.16 | -0.47 | 0.72 | $8.60x10^5$ | $3.03x10^6$ |
| 7 | 31.92 | -0.85 | -0.68 | 0.44 | $4.62x10^5$ | $8.19x10^5$ |
| 8 | 35.55 | 0.38 | -0.44 | 0.51 | $3.23x10^5$ | $6.62x10^5$ |
| 9 | 58.23 | 0.16 | -0.41 | 0.73 | $3.47x10^5$ | $1.29x10^6$ |
| 10 | 122.71 | -0.28 | -0.60 | 0.71 | $1.75x10^6$ | $5.97x10^6$ |
| 11 | 30.78 | -0.15 | -0.66 | 0.72 | $1.28x10^5$ | $4.53x10^5$ |
| 12 | 21.62 | 0.02 | -0.89 | 0.35 | $3.20x10^5$ | $4.96x10^5$ |
| 13 | 72.60 | 0.17 | -0.50 | 0.76 | $4.88x10^5$ | $2.00x10^6$ |
| 14 | 52.90 | -2.14 | 0.71 | 0.27 | $2.55x10^6$ | $3.46x10^6$ |
| 15 | 47.46 | -0.10 | -0.26 | 0.52 | $5.67x10^5$ | $1.18x10^6$ |
| 16 | 51.53 | -0.06 | -0.86 | 0.58 | $6.47x10^5$ | $1.55x10^6$ |
| 17 | 76.58 | -0.38 | -0.39 | 0.62 | $9.43x10^5$ | $2.46x10^6$ |
| 18 | 41.05 | -0.21 | -0.65 | 0.55 | $4.76x10^5$ | $1.06x10^6$ |
| 19 | 21.18 | 0.01 | -0.26 | 0.39 | $2.69x10^5$ | $4.41x10^5$ |
| 20 | 28.35 | -0.44 | 0.40 | 0.61 | $2.10x10^5$ | $5.42x10^5$ |
| 21 | 49.87 | 0.46 | -0.38 | 0.51 | $7.73x10^5$ | $1.58x10^6$ |
| 22 | 63.68 | -0.70 | 0.14 | 0.57 | $9.61x10^5$ | $2.23x10^6$ |
| 23 | 52.40 | -0.74 | -0.54 | 0.43 | $1.01x10^6$ | $1.77x10^6$ |
| **Mean (s.e.m.)** | **53.93 (5.74)** | **-0.29 (0.17)** | **-0.41 (0.08)** | **0.58 (0.03)** | **$6.95x10^5$ ($1.21x10^5$)** | **$1.77x10^6$ ($3.16x10^5$)** |

**Supplementary Table 3:** Parameters corresponding to the best sinusoidal model for each participant's 'breathing phase ~ interbeat interval' curve based on the last three R-peaks from the stimulus anticipation periods in each trial. (A: amplitude, B: vertical offset, φ= phase shift, $R^2$: R-squared (coefficient of determination), $SS_{res}$: residual sum of squares, $SS_{tot}$: total sum of squares)

| Subj. ID | A | B | ϕ | $R^2$ | $SS_{res}$ | $SS_{tot}$ |
|---|---|---|---|---|---|---|
| 1 | 22.86 | 3.52 | -0.23 | 0.49 | $3.13x10^5$ | $6.09x10^5$ |
| 2 | 96.24 | 21.74 | -0.43 | 0.50 | $6.10x10^6$ | $1.22x10^7$ |
| 3 | 53.21 | 12.50 | -0.33 | 0.70 | $7.21x10^5$ | $2.44x10^6$ |
| 4 | 58.10 | 33.57 | -0.45 | 0.44 | $2.44x10^6$ | $4.37x10^6$ |
| 5 | 32.87 | 16.19 | -0.79 | 0.46 | $9.67x10^5$ | $1.78x10^6$ |
| 6 | 106.80 | 15.46 | -0.56 | 0.66 | $4.05x10^6$ | $1.19x10^7$ |
| 7 | 20.13 | 3.94 | -0.82 | 0.31 | $6.59x10^5$ | $9.51x10^5$ |
| 8 | 30.37 | 3.50 | -0.79 | 0.40 | $8.99x10^5$ | $1.50x10^6$ |
| 9 | 57.37 | 9.30 | -0.49 | 0.66 | $1.16x10^6$ | $3.46x10^6$ |
| 10 | 95.69 | 23.95 | -0.56 | 0.48 | $7.59x10^6$ | $1.47x10^7$ |
| 11 | 22.33 | 4.15 | -0.69 | 0.47 | $4.45x10^5$ | $8.46x10^5$ |
| 12 | 20.73 | 12.68 | -0.86 | 0.18 | $1.45x10^6$ | $1.78x10^6$ |
| 13 | 43.28 | 17.57 | -0.52 | 0.47 | $1.22x10^6$ | $2.33x10^6$ |
| 14 | 55.91 | 13.60 | -0.70 | 0.44 | $2.01x10^6$ | $3.58x10^6$ |
| 15 | 44.80 | 3.35 | -0.41 | 0.49 | $1.64x10^6$ | $3.20x10^6$ |
| 16 | 48.32 | 13.95 | -0.83 | 0.47 | $1.54x10^6$ | $2.91x10^6$ |
| 17 | 43.07 | 7.20 | -0.10 | 0.39 | $2.04x10^6$ | $3.37x10^6$ |
| 18 | 51.08 | 7.51 | -0.65 | 0.51 | $1.59x10^6$ | $3.24x10^6$ |
| 19 | 17.07 | 1.40 | -0.69 | 0.27 | $4.59x10^5$ | $6.32x10^5$ |
| 20 | 47.49 | 10.69 | 0.05 | 0.72 | $6.09x10^5$ | $2.17x10^6$ |
| 21 | 55.85 | 13.22 | -0.35 | 0.42 | $3.31x10^6$ | $5.70x10^6$ |
| 22 | 83.89 | 8.33 | -0.11 | 0.59 | $2.69x10^6$ | $6.51x10^6$ |
| 23 | 51.15 | 7.39 | -0.52 | 0.54 | $1.28x10^6$ | $2.77x10^6$ |
| **Mean (s.e.m.)** | **50.37 (5.25)** | **11.51 (1.61)** | **-0.52 (0.05)** | **0.48 (0.03)** | **$1.97x10^6$ ($3.79x10^5$)** | **$4.04x10^6$ ($8.01x10^5$)** |

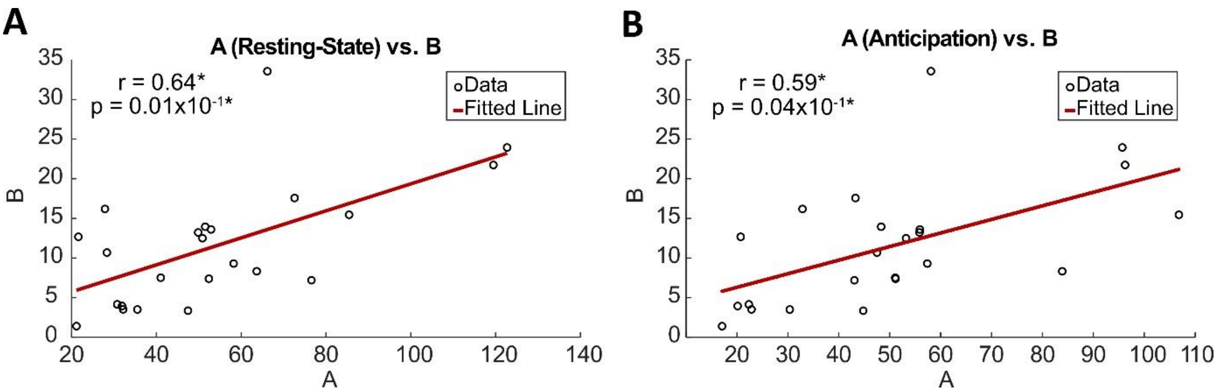


**Supplementary Figure 1: Linear correlations between vertical offset (B) and RSA. A)** RSA during the 10-minute resting-state, **B)** average RSA during stimulus anticipation. Values of A correspond to those shown in Supplementary Tables 2 and 3, respectively; whereas values of B are from the stimulus anticipation period on both graphs (*: r and p-values are derived from robust linear regression).

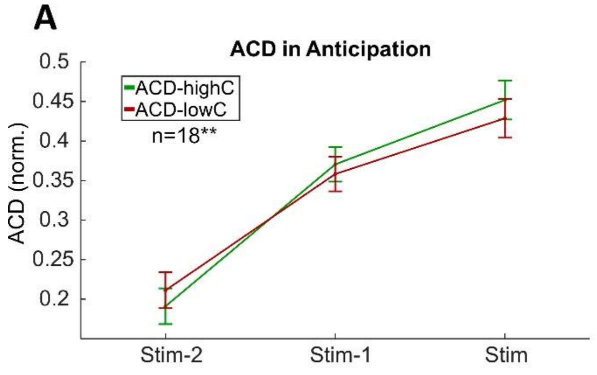


| ACD | SumSq | DF | MeanSq | F | pGG |
|---|---|---|---|---|---|
| Confidence | $6.91 \times 10^{-4}$ | 1 | $6.91 \times 10^{-4}$ | 0.33 | 0.58 |
| Time | 0.92 | 2 | 0.54 | 14.39 | $5.52 \times 10^{-4}$ |
| Interaction | 0.01 | 2 | $0.05 \times 10^{-1}$ | 2.09 | 0.14 |

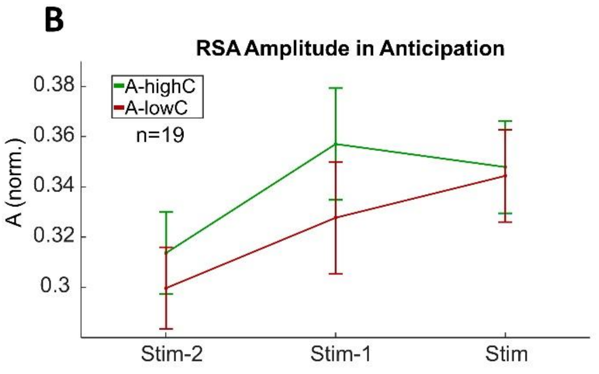


| RSA Amplitude (A) | SumSq | DF | MeanSq | F | pGG |
|---|---|---|---|---|---|
| Confidence | 0.01 | 1 | 0.01 | 2.21 | 0.15 |
| Time | 0.04 | 2 | 0.02 | 4.67 | 0.02 |
| Interaction | $3.21 \times 10^{-3}$ | 2 | $1.60 \times 10^{-3}$ | 2.03 | 0.16 |

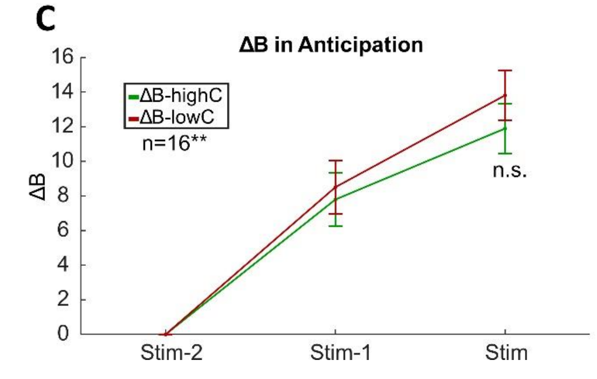


**Supplementary Figure 2: RSA and respiratory phase-independent cardiac deceleration in High- and Low-confidence trials. A)** Total cardiac deceleration, **B)** Estimated RSA amplitude during stimulus anticipation in High-confidence and Low-confidence trials ('highC' and 'lowC' in figure legends, respectively). **C)** Progress of the vertical offset parameter within stimulus anticipation. The time point of "R:stim-2" heartbeat was taken as reference (n.s.: p>0.05; ** in **A** and **C**: in addition to the 4 participants that were excluded from confidence analysis due to outlier confidence ratings, additional participant(s) had to be excluded due to outlier cardiorespiratory parameter values). Error bars correspond to within-participant 95% confidence intervals. Median confidence ratings of 4 subjects were 100%, therefore they could not be included in the confidence analyses based on median-splitting the ratings to categorize the trials as low- and high-confidence trials.

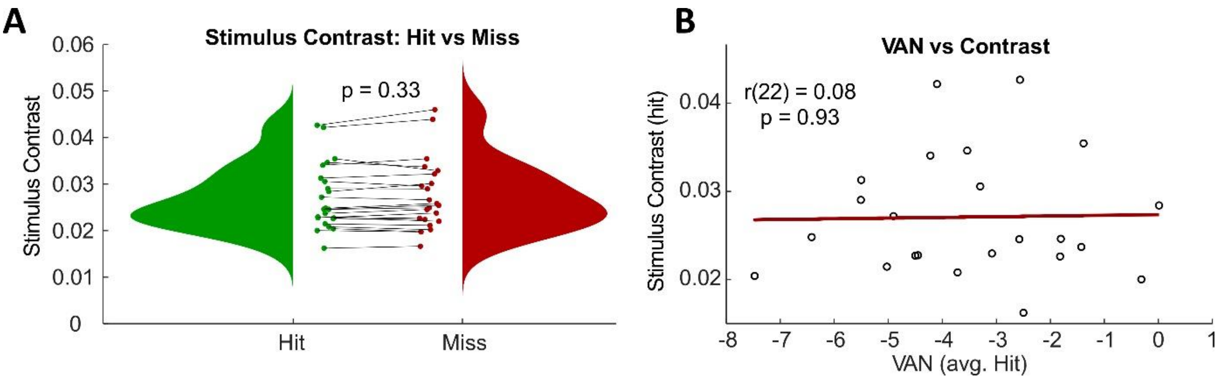


**Supplementary Figure 3: Controlling for the potential effect of physical stimulus contrast on Visual Awareness Negativity (VAN). A)** Comparison of average stimulus contrasts shown to each participant after downsampling Hit trials to match the Miss trials in average contrast values (p-value corresponds to Wilcoxon signed-rank test). **B)** VAN values do not reflect the average stimulus contrast shown to the participants (r: Spearman's correlation coefficient, p: p-value for the significance of the correlation based on robust linear fitting).
